# Genotype-Phenotype Relations of the von Hippel-Lindau Tumor Suppressor Inferred from a Large-Scale Analysis of Disease Mutations and Interactors

**DOI:** 10.1101/405845

**Authors:** Giovanni Minervini, Federica Quaglia, Francesco Tabaro, Silvio C.E. Tosatto

## Abstract

Familiar cancers represent a privileged point of view for studying the complex cellular events inducing tumor transformation. Von Hippel-Lindau syndrome, a familiar predisposition to develop cancer is a clear example. Here, we present our efforts to decipher the role of von Hippel-Lindau tumor suppressor protein (pVHL) in cancer insurgence. We collected high quality information about both pVHL mutations and interactors to investigate the association between patient phenotypes, mutated protein surface and impaired interactions. Our data suggest that different phenotypes correlate with localized perturbations of the pVHL structure, with specific cell functions associated to different protein surfaces. We propose five different pVHL interfaces to be selectively involved in modulating proteins regulating gene expression, protein homeostasis as well as to address extracellular matrix (ECM) and ciliogenesis associated functions. These data were used to drive molecular docking of pVHL with its interactors and guide Petri net simulations of the most promising alterations. We predict that disruption of pVHL association with certain interactors can trigger tumor transformation, inducing metabolism imbalance and ECM remodeling. Collectively taken, our findings provide novel insights into VHL-associated tumorigenesis. This highly integrated *in silico* approach may help elucidate novel treatment paradigms for VHL disease.

**Author summary:** Cancer is generally caused by a series of mutations accumulating over time in a healthy tissue, which becomes re-programmed to proliferate at the expense of the hosting organism. This process is difficult to follow and understand as events in a multitude of different genes can lead to similar outcomes without apparent cause. The von Hippel-Lindau (VHL) tumor suppressor is one of the few genes harboring a familiar cancer syndrome, i.e. VHL mutations are known to cause a predictable series of events leading cancer in the kidneys and a few selected other tissues. This article describes a large-scale analysis to relate known VHL mutations to specific cancer pathways by looking at the molecular interactions. Different cancer types appear to be caused by mutations changing the surface of specific parts of the VHL protein. By looking at the VHL interactors involved, it is therefore possible to identify other candidate genes for mutations leading to very similar cancer types.

## Introduction

Familial cancers are rare, accounting for about 5-10% of all cancers (1-3) and generally characterized by inherited inactivation of important tumor suppressors. Inherited tumors represent a valuable source of information about the mechanisms driving cancerogenesis. These cancers are associated to mutations of known genes, allowing the formulation of clear genotype-phenotype correlations in many cases. The von Hippel-Lindau (VHL) syndrome is a familial disorder characterized by a predisposition to develop several different benign and malignant tumors, such as retinal- and cerebellar-hemangioblastoma, pheochromocytoma, paraganglioma, nonfunctioning pancreatic neuroendocrine tumors (pNETs) and renal cell carcinoma (RCC) (4-6). VHL syndrome arises from pathogenic inactivation of the von Hippel-Lindau gene located on chromosome three (4), which codes for the homonymous pVHL protein. pVHL is mainly known to act as substrate recognition component of a protein complex (7) formed together with elongin-B, elongin-C and cullin-2 (VCB), possessing ubiquitin ligase E3 activity towards the HIF-1α transcription factor (8). Association between pVHL and HIF-1α is oxygen-dependent and triggered through hydroxylation by the PHD (prolyl hydroxylase domain containing) enzyme family of two HIF-1α proline residues (7-10). PHD activity is itself inhibited under hypoxia, allowing HIF-1α to escape degradation and translocate to the nucleus where it activates hypoxia-dependent target genes. Clinically, VHL disease is classified as Type 1 or Type 2 depending on clinical manifestations in patients (11). Type 1 includes patients harboring either truncating mutations or deletions yielding a dysfunctional pVHL presenting a wide spectrum of different cancers but lacking pheochromocytoma. Type 2 is more genetically divergent, characterized by missense mutations and includes patients developing pheochromocytoma (11). Type 2A presents pheochromocytoma and other typical VHL manifestations (e.g. cysts) except RCC (renal cell carcinoma). Type 2B covers almost the entire spectrum of VHL manifestations including aggressive RCC, while type 2C only develops isolated pheochromocytoma. Although routinely used for the initial assessment of VHL patients, this classification can generate ambiguous assignments. Clinical VHL manifestation is frequently variable, with different phenotypes in different families or even in the same family (12). Several different functions were attributed to pVHL in addition to its role in HIF-1α degradation in light of these variable phenotypes. pVHL has been reported to associate and promote p53 stabilization upon MDM2 inhibition (13), mediate assembly and regulation of the extracellular matrix (ECM) (14-17), regulate cell senescence (18) and apoptosis (19,20) as well as playing a role in regulating oxidative stress gene transcription response (21). pVHL is also known to harbor at least three different binding surfaces (22) which are thought to be involved in multiple protein-protein interactions. While hundreds of different pVHL protein-protein associations are described (23), whether and how these HIF-independent pVHL contribute to VHL tumorigenesis is largely unknown. Here, we present our efforts in deciphering pVHL function. A thorough investigation of pVHL interactors and binding surfaces was coupled with manual curation of pathogenic mutations. Mutations predicted to impair specific pVHL functions were associated with the corresponding VHL phenotype, while a list of affected pathways was constructed for each phenotype. Our analysis shows that the different phenotypes described in VHL patients correlate with specific structural pVHL perturbations, showing how pVHL interfaces to correlate differentially with specific phenotypes. Our data also show that some HIF1-independent functions of pVHL can be attributed to specific pVHL regions.

## Results

Although the best known pVHL function is its role in degrading the HIF-1α transcription factor (4), a number of HIF-independent functions have also been reported (24,25). We previously proposed three distinct pVHL interaction interfaces (A, B, and C) (22), engaging in exclusive interactions with different interactors and partially explaining the pVHL binding plasticity. Based on the available pVHL 3D structures, interface A is important for VCB complex formation (26). Interface B forms the HIF-1α binding-site (7), while interface C participates in the cullin2 interaction (26). A further interface is defined by an accessory N-terminal tail (surface D) present only in the pVHL30 isoform (23). As the recently developed VHLdb database (23) collects pVHL interactors and mutations, here we present an analysis of this data to shed light on pVHL functions. VHLdb missense mutations were mapped on the pVHL structure to highlight specific interaction surfaces. Deletions of vast pVHL regions were excluded as they yield a clearly dysfunctional protein. A detailed investigation of pVHL interactors was carried out to assess their ability to explain different VHL phenotypes. Collectively, our analysis covers 59 pVHL interactors and 379 different mutations from 2,123 VHL patients described in 1,597 case reports from 143 scientific papers. These data were used to list the most affected pVHL interactors in VHL disease and their effects were simulated using a Petri net (27).

### pVHL mutation hot spots

As pVHL mutations appear to indiscriminately affect the entire protein surface, we wondered whether some mutations were more frequent in patients. We started analyzing the pVHL mutation distribution and identified 18 frequently mutated pVHL positions (>30 occurrences) forming mutation hotspots (Fig. 1). Most residues present multiple amino acids substitutions, e.g. 7 different variants are described for Asn78, and localize in both alpha and beta pVHL domains. Imposing a cutoff >30 patients, ten different single mutations were found to be the most recurrent (incidence > 33%) (Figure 1 and S1 Table). All associate with malignant RCC and other typical VHL manifestations, while only one is found in patients with polycythemia. Arg167 is the most frequently mutated pVHL residue (338 patients), with the two main variants p.Arg167Trp and p.Arg167Gln. This residue localizes on surface A, which is involved in VCB complex formation. Inspection of complexes formed by pVHL, Elongin-B and -C and cullin-2 with RING(28) shows the residue to be located on the inner side of pVHL α-helix 1, directly interacting with Elongin-C (Figure 2). Arg167 forms salt bridges with Glu160 and Asp126, playing a structural role in maintaining the correct α-domain fold. Intriguingly, the phenotypes described for the two variants are slightly different. Both associate with RCC, pheochromocytoma, hemangioblastoma, paraganglioma and cyst formation. Patients harboring p.Asp167Trp are also found to develop epididymal/ovarian cystadenoma (1 occurrence), while presence of endolymphatic sac tumor is reported only for p.Asp167Gln (2 occurrences). Despite the exiguous number of reported patients, this finding suggests that the two changes may support different tissue-specific pVHL functional impairment. The p.Arg161Gln mutation destabilizes the interaction between pVHL and Elongin-C. This arginine forms a salt-bridge with Elongin-C Glu92 and substitution with glutamine impairs this interaction, weakening VCB complex formation. Collectively, our analysis shows the most frequent mutations affecting surface A to exert pathogenicity by directly impairing VCB complex formation rather than compromising the pVHL/HIF-1α interaction. Conversely, mutation p.Tyr98His on surface B exerts its pathogenic effect by directly disrupting pVHL association with HIF-1α. Tyr98 plays a predominant role in pVHL substrate recognition, forming a hydrogen bond with the HIF1-α Pro564 backbone. Of note, this specific HIF1-α proline is crucial for pVHL interaction upon hydroxylation (7). Remaining frequent mutations localizing on surface B (p.Ser65Leu, p.Asn78Ser, Pro81Ser) are predicted to play a role in destabilizing the β-domain. Structural investigation shows these residues mostly face the inner layer of the β-domain and are not directly in contact with conventional pVHL interactors, i.e. Elongin-B and -C, at least based on the currently available crystal structures. Similarly, mutation p.Ala149Ser affects surface C and is localized on the seventh strand of the β-domain. Ala149 contributes to the pVHL hydrophobic core, forming van der Waals interactions with Val74, Phe76, Phe119, Val130 and Phe136. Substitution with serine may drive pVHL fold destabilization and VHL syndrome insurgence. Finally, p.Arg200Trp is the only frequent mutation in the pVHL C-terminal tail and mostly associated to polycythemia (S1 Table). It disrupts a salt bridge between Arg200 and Glu134 which helps the α-domain to fold correctly.

**Figure 1:**
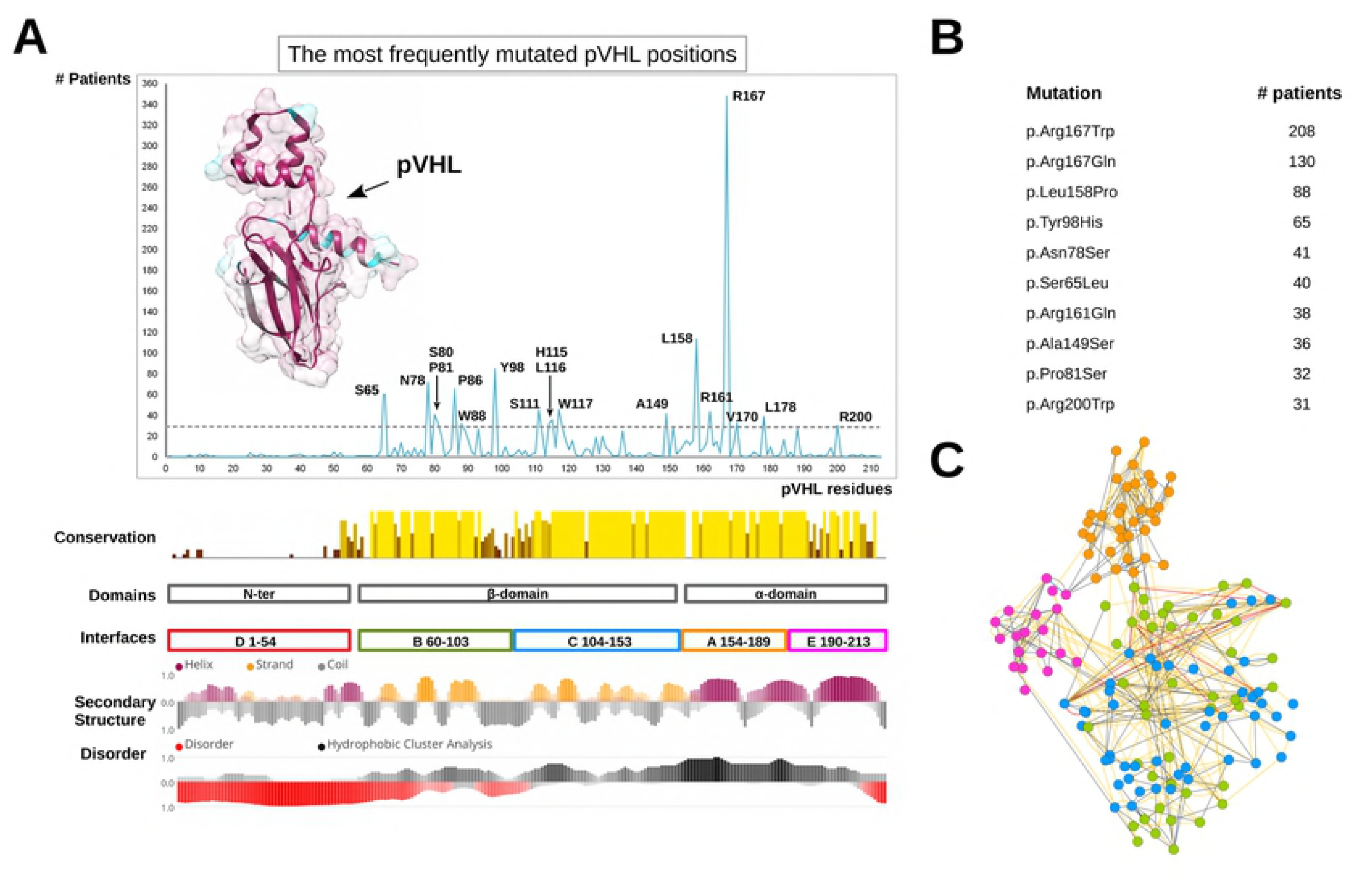
The most frequently mutated pVHL positions. (A) pVHL mutations are grouped and collected based on their occurrence among VHL patients. The pVHL 3D structure is presented as cartoon and colored by sequence conservation. Interfaces definition, secondary structure and disorder content are presented below. (B) The ten most frequent pVHL mutation affecting >30 patients. (C) RING representation of pVHL residues interaction network, colored by secondary structure. Edges represent connection among residues (nodes), with red representing π-π stack interaction, blue salt bridges while yellow and grey for van der Waals and hydrogen bonds respectively.

**Figure 2:**
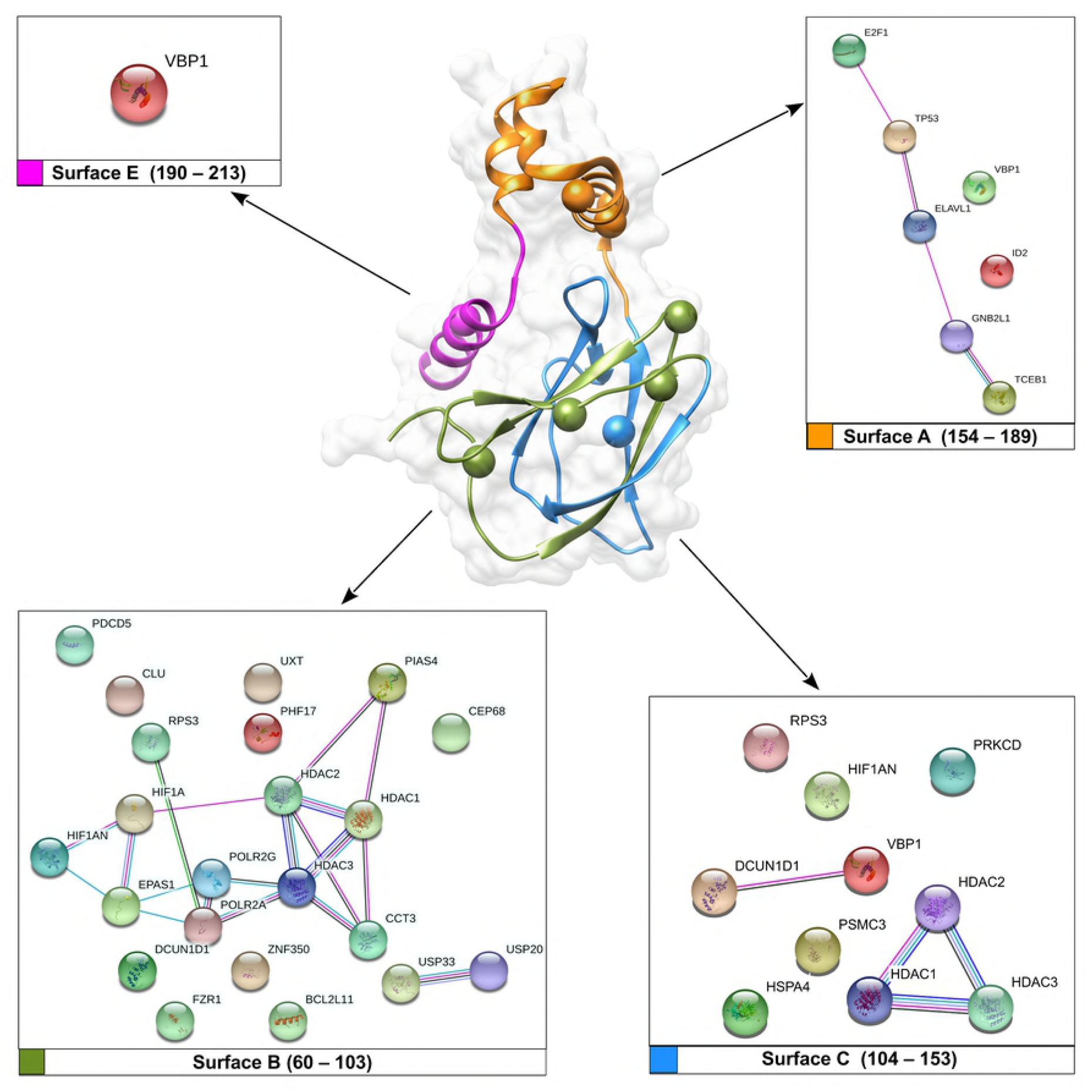
Network of pVHL interacting proteins and associated pVHL surfaces. Manually curated pVHL interactors from VHLdb are represented as colored sphere with connecting edges representing physical interaction among them.

### Protein networks impaired by the most frequent pVHL mutations

Our preliminary mapping of the most recurring pVHL mutations provides some useful hints about their pathogenic effect. Such an interpretation however fails to explain the variability of phenotypes observed in VHL patients (S2 Table). A possible interpretation is that these amino acid changes disrupt pVHL association with interactors other than Elongin-C, Elongin-B and HIF-1α. To address this issue, we extracted from VHLdb interactors known to specifically bind pVHL regions corresponding to the most frequent mutations (S7 File). These were used to generate protein association networks describing the putatively compromised cellular processes (Fig. 3). The surface A interaction network collects eight proteins mainly involved in DNA/RNA processing. pVHL loss is frequently associated to genomic instability in renal cancer (29). In particular, pVHL null cells display reduced activation of p53-mediated apoptotic response (30), as well as abnormal cell-cycle arrest upon DNA damage, with normal response observed after pVHL restoration. We searched for highly interconnected regions of the network around surface A using MCODE (31) to better characterize these observations. Three main clusters representing macromolecular protein complexes were found (S8 file). The first cluster is formed by five proteins belonging to the prefoldin protein family, a group of chaperon forming folding complexes (32). As pVHL is thought to interact with more than four-hundred different interactors (23), different prefoldin proteins may assist formation of multiple pVHL-driven protein complexes. Surface A mutations may interfere with complex assembly, promoting abnormal cell behavior. The second cluster accounts for proteins directly involved in p53 activation, supporting a pVHL role in this specific cellular function. The last cluster for surface A includes Elongin-C and Elongin-B, confirming the role of this interface in VCB complex assembly. Of note, this cluster also includes SOCS3 (Suppressor of cytokine signaling 3), a protein acting as negative regulator of cytokine signal transduction. SOCS3 binds pVHL to form a heterodimeric E3 ligase complex targeting the JAK2 kinase for degradation (33). This interaction was proposed to play a role in Chuvash polycythemia insurgence (33), a familiar polycythemia form caused by pVHL mutations. Collectively, our findings suggest that interface A is involved in formation of multiple protein complexes. Mutations affecting this region may result in abnormal gene transcription as well as deregulation of apoptotic response and signaling. Mutations on surface B can compromise pVHL association with 23 different interactors. Surface B contains the HIF-1α binding site and is also the binding interface of multiple proteins involved in regulating the pVHL E3-ligase activity. The two deubiquitinating enzymes USP33 and USP20 are clear examples. This specific enzyme class is involved in regulating multiple pathways by modulating protein degradation. USP33 and USP20 in particular play a relevant role in beta-adrenergic receptor (ADRB2) homeostasis. Upon prolonged agonist stimulation, they constitutively bind ADRB2 and inhibit its lysosomal trafficking (34). The pVHL interaction with at least three different histone deacetylases was also found putatively compromised. In addition to being an E3 ligase component, pVHL binds histone deacetylases to form a heterodimeric complex, acting as a transcriptional co-repressor to inhibit the HIF-1α trans-activation function (35). Unexpectedly, our network analysis of surface B interactors shows only one relevant cluster accounting for ten ribosomal proteins. Recently, pVHL was proposed to inhibit both ribosome biogenesis and protein synthesis by inducing nuclear retention of pre-40S ribosomal subunits (36). The biological meaning of this interaction is however far from understood. Taken together, our findings suggest surface B to be mainly involved in protein degradation pathways. Based on these data, we suggest that the pathogenic assessment of mutations affecting this area should also include other putative pVHL hydroxylated-substrates beyond the sole HIF-1/2α, e.g. SPRY2 (37), ADRB2 (38), EPOR (39). Eleven interactors are found to be affected by surface C mutations. Cluster analysis suggests this interface also to play a role in ribosome biogenesis and protein synthesis, as already observed for surface B. We found a second cluster collecting members of the histone-deacetylation protein family, i.e. HDAC1-3. This suggests interface C to also have an important role in transcriptional regulation and cell cycle progression. Finally, a single cluster collecting proteins involved in matrix organization, ciliogenesis and BBsome assembly (Bardet-Biedl syndrome, an octameric protein complex required for ciliogenesis and centriolar function (40)) was found to specifically interact with the pVHL C-terminus, suggesting this region may form a further pVHL interaction interface (S8 file).

**Figure 3:**
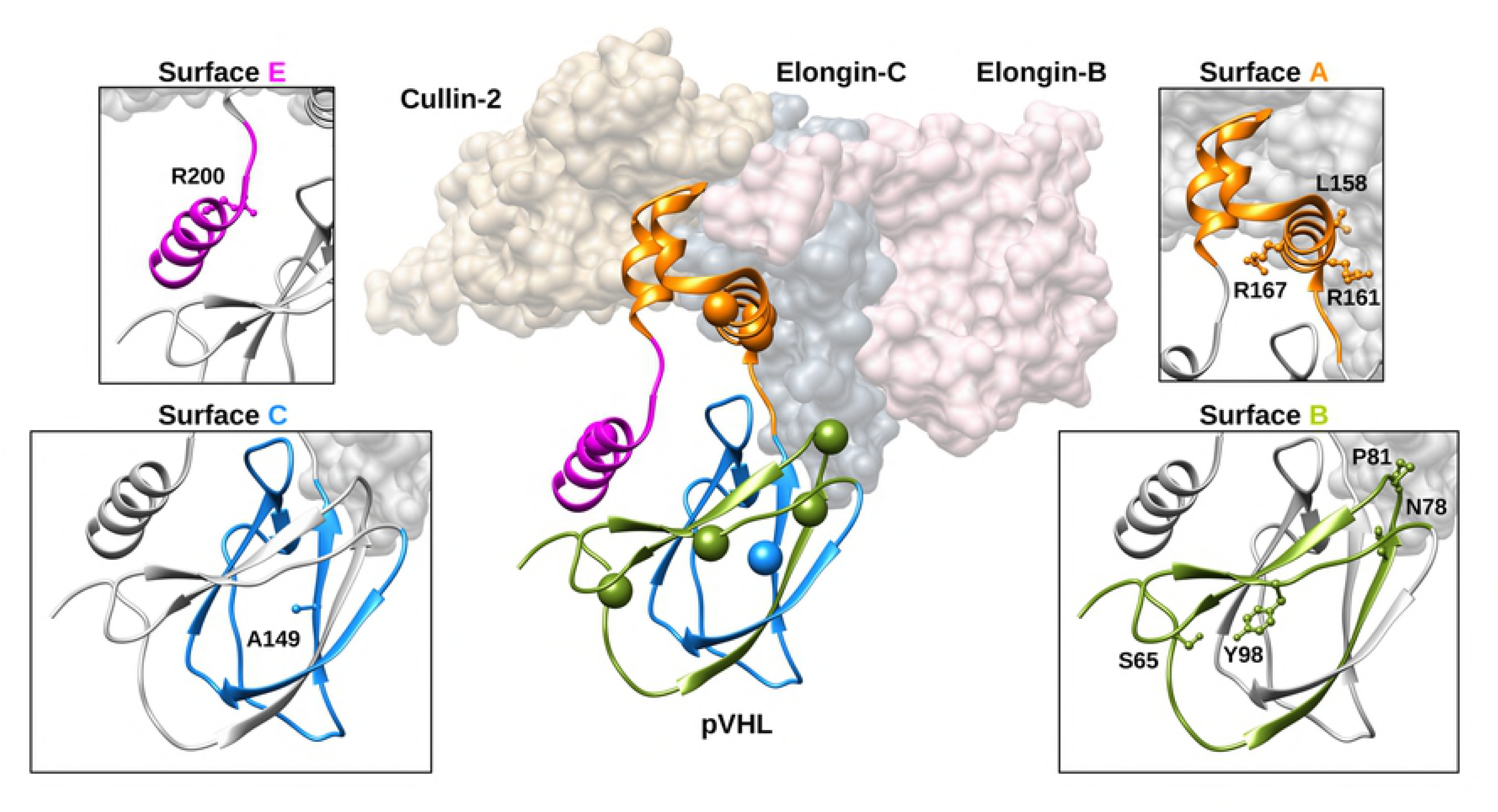
Overview of the VCB complex. Elongin-B and -C and Cullin2 are represented as plain surface and colored in pink, dark grey and brown, respectively. Cartoon represents pVHL, with the ten most mutated residues presented as spheres. Zooms show different pVHL binding interfaces colored according residues forming the interface. In particular, orange is for surface A, while green and blue are for surfaces B and C respectively. C-terminal region (surface E) is presented in magenta.

### Distribution of pVHL mutations and associated phenotypes

The current classification describes two main types of VHL disease based on their propensity to develop pheochromocytoma (11), Type 1 (low risk) and 2 (high risk), respectively. Three sub-types are further proposed for Type 2 to appraise the risk of developing renal carcinomas. These differences can be interpreted as resulting from impairment of different protein-protein interactions. We wondered whether the position of mutations on pVHL surfaces may associate with specific disease manifestations. We retrieved 1,597 mutations affecting different pVHL binding interfaces (22,23) and isolated 726 amino acid variations associated to VHL phenotype, e.g. only substitutions described as yielding RCC (S3 Table). We focus on mutations affecting single residues on each binding surface, to isolate pVHL areas that can support unknown functions or binding motifs. Unsurprisingly, our analysis shows mutations to distribute over the entire protein surface. However, differences in localization were observed (Fig. 4). Normalizing the number of mutated positions over the number of residues forming each interface shows surface B to present the highest number of mutated positions (39/43), indicating that 91% of residues forming this interface are targeted by VHL mutations. Surface C (43/49; 88%) and surface A (28/35; 80%) have slightly fewer mutations, followed by the C-terminal tail (15/23; 65%) and interface D (27/59; 56%). Surfaces B and C form together the β-domain of pVHL, which includes the HIF-1α binding site. These preliminary findings can be easily explained accounting for the specific domain function. Surface A includes the pVHL α-domain needed to sustain interaction with Elongin-B and -C to form the VCB complex. The meaning of mutations affecting both surface D and the protein C-terminus is more difficult to address. Surface D is an accessory acidic tail present only in the pVHL30 isoform (41). A pVHL30-specific role in regulating p14ARF tumor-suppressor activity has been proposed (42), however this function is currently debated. We investigated whether phenotypes associated with these mutations can be used to draw surface/phenotype correlations. Our analysis shows that the substitution of residues forming interfaces B and C yields similar phenotypes (Fig. 4). Renal disease, globally including ccRCC, RCC and renal cysts, is on average the main manifestation associated with these two interfaces. A similar tendency is also observed for hemangioblastoma (both cerebellar- and retinal-forms) and pancreatic lesions (cysts and tumors). Pheochromocytoma is slightly more frequent when mutations affect surface B (17.5%) than surface C (15.3%). Impairment of surface C is also associated to other minor phenotypes which are virtually absent in surface B, e.g. colorectal cancer and endolymphatic sac tumor. In the same way, surface A mutations are associated with almost the entire spectrum of VHL phenotypes (Figure 4). Renal disease (27.7%), pheochromocytoma (20.5%) and hemangioblastoma (34.4%) are similarly distributed. Minor manifestations are also present and uniformly distributed, even though accounting for smaller numbers (< 3.5%). Conversely, mutations affecting surface D and the C-terminal tail describe a different scenario. Surface D mutations, which are only present in the pVHL30 isoform, are mostly associated with renal disease (75.6%). Of note, cerebellar-hemangioblastoma (6.7%) is the only hemangioblastoma subtype described for mutations affecting this interface. This suggests that pVHL30 may play a cerebellar-specific role which has no equivalent in retinal tissue. Finally, C-terminal mutations are mainly characterized by renal syndrome (40.7%) and pheochromocytoma (18.5%). Intriguingly, mutations associated to non-canonical VHL manifestations, such as polycythemia (14.8%), colorectal cancer (14.8%) and glial tumor (7.4%) seem mostly to derive from mutations of this interface. Structural investigation suggests that changes in this region should neither impair HIF-1α nor VCB complex formation. These findings collectively suggest that the C-terminal region may play a functional role in other HIF-independent pVHL functions. The analysis confirms that the three main interfaces (A-C) present no statistically significant difference in tumor type association. Conversely, the different correlations observed for the two pVHL tails are statistically significant (p-values <0.05, S3 Table). Analyzing the data presented so far from a disease point of view, we observe that renal disease, pheochromocytoma and pancreatic insults derive from mutations affecting all pVHL interfaces. Similarly, pheochromocytoma behavior suggests that both phenotypes are associated with general pVHL impairment. Instead, retinal-hemangioblastoma arises from mutations limited to the three main interfaces (A-C). The cerebellar subtype also seems to include mutations on interface D present only in the pVHL30 isoform. It will be very interesting to investigate whether these differences can be explained with a functional specialization of pVHL30 in cerebellar tissues.

**Figure 4:**
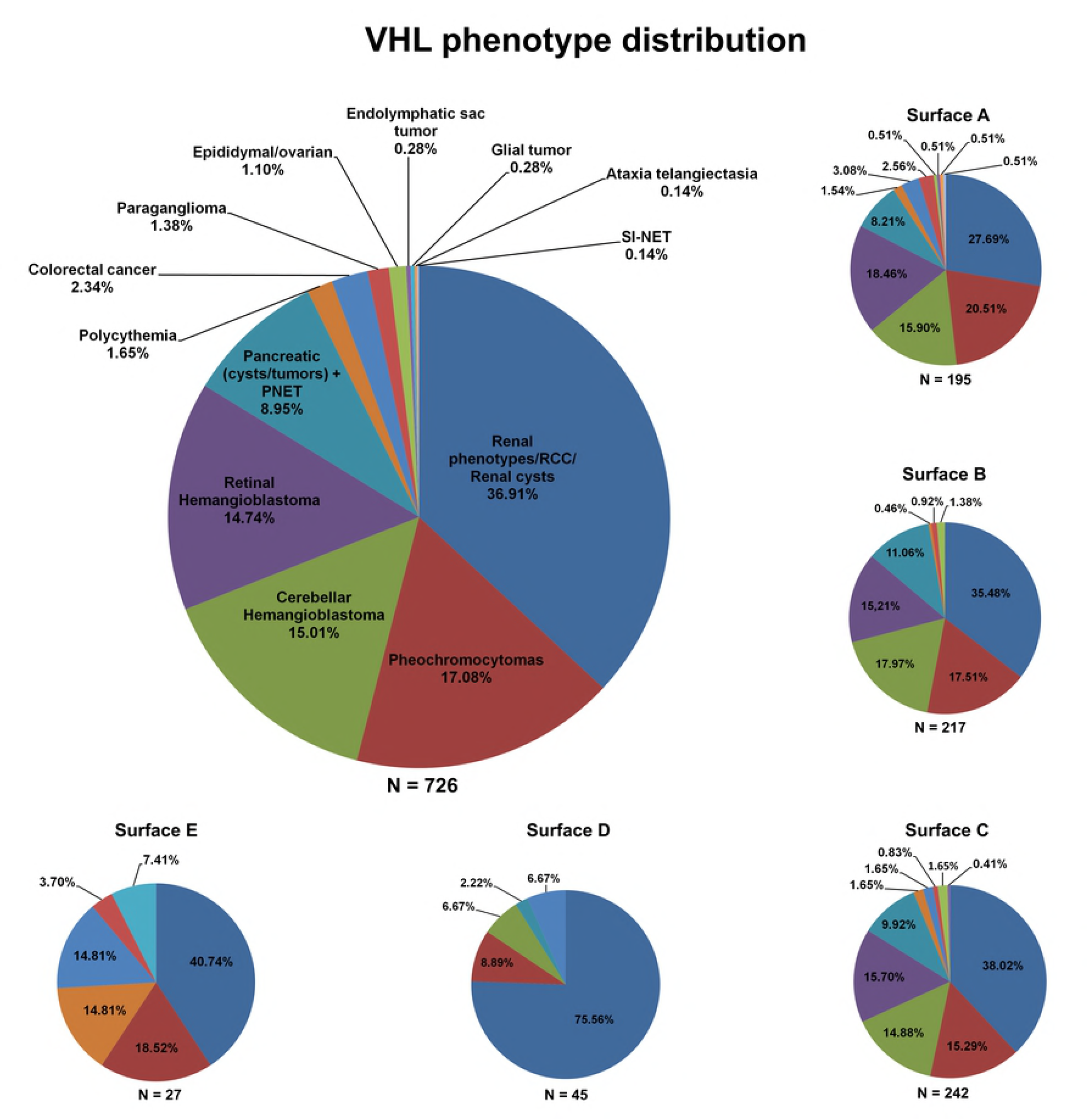
Distribution analysis of pVHL mutations and related phenotypes. The large circle shows the percentage of mutations for which a specific phenotype is reported from patients. The same distribution is reported for each pVHL surface (small circles). HB = hemangioblastoma; PNET =pancreatic neuroendocrine tumor; RCC = renal cell carcinoma; SI-NET = small intestine neuroendocrine tumor.

### pVHL interactors and disease states

VHL disease is characterized by slow progression coupled to a plethora of different symptoms. Detailed molecular data for 56 pVHL interactors involves pVHL surfaces reported to engage in multiple protein-protein interactions (23,42,43). We selected VHLdb single point mutations reported to associate with a single specific VHL manifestation to suggest binding interfaces correlated with specific pVHL functions (S4 Table). Unsurprisingly, we observed that many mutations included in this reduced subset affect regions involved in protein-protein complex formation. This finding suggests a single amino acid substitution can impair formation of multiple associations. On the other hand, it makes retrieving which specific interaction is actually weakened or damaged by the mutation more difficult. We decided to isolate only pVHL interactors unaffected by the mutation associated to a specific phenotype to reduce incoherence and lower the risk of redundancy, generating a set of negative snapshots describing pVHL interactions and pathways not directly compromised by a specific mutation (Table 1). Our data show that mutations promoting RCC, and more in general renal disease, can interest virtually all considered interactors. This is coherent with an almost complete functional inactivation of pVHL, in particular for functions connected with hypoxia response. Notably, post translational modification (PTM) sites present on pVHL are frequently mutated in RCC (and renal disease). We found mutations in PTM sites recognized by AURKA, CSNK2A1, CHEK2, GSK3B, as well as NEDD8. The latter, a ubiquitin-like molecule, is thought to activate a pVHL functional switch promoting detachment from the VCB complex and allowing association with fibronectin (44). Inactivation of either phosphorylation or neddylation sites may trigger RCC development. Similarly, mutations only associated with pheochromocytoma severely affect the DGKZ, PRKCI, PRKCD and PRKCZ kinase binding interfaces, further suggesting a link between hypoxia sensing and phosphorylation-mediated signaling. Mutations associated only to this neuroendocrine tumor affect the pVHL association with almost the entire subset of interactors considered. Notably, associations with AKT1 and HSPA4 are not impaired in pheochromocytoma. AKT1 is known to promote survival and proliferation of various cancers (45). Its interaction with pVHL is proline-hydroxylation-dependent and yields complete inhibition of AKT1 activity (46). Our finding indicates that cells harboring these mutations may retain AKT1 inhibition and suggest that proliferative pathways modulated by pVHL/AKT1 association are not compromised. The retained HSPA4 interaction seems to go in exactly the same direction. HSPA4, also known as HSP70RY (47), is a member of the HSP70 chaperone family proposed to rescue premature degradation of pVHL mutants, inducing their stabilization and halting tumor progression (48). Collectively, these data are in agreement with the slow progression rate reported for pheochromocytomas (49) and describe this cancer to arise only from partial pVHL inactivation. Mutations associated with both cerebellar- and retinal-hemangioblastoma localize in regions not affecting the pVHL interaction with transcription factors (TP53, ELAVL1) and regulators (ID2, E2F1, ELOC). Although pVHL is known to interact with two RNA polymerase II subunits, RBP1 (50) and RBP7 (51), its role in regulating gene transcription is debated (25). According to our data, cell damage promoting hemangioblastoma should be ascribed to inactivation of HIF-dependent functions. Conversely, pVHL mutations associated to colorectal cancer do not affect association with HIF-1α, TP53, or proteins involved in extracellular matrix (ECM) formation and turnover, such as TUBB, TUBA1A, TUBA4A, KIF3A and KIF3B. Finally, a single mutation is observed as never inducing both paraganglioma, a tumor originating from paraganglia in chromaffin-negative glomus cells, and cystadenoma, a tumor of epithelial tissue with glandular origin.

**Table 1:**
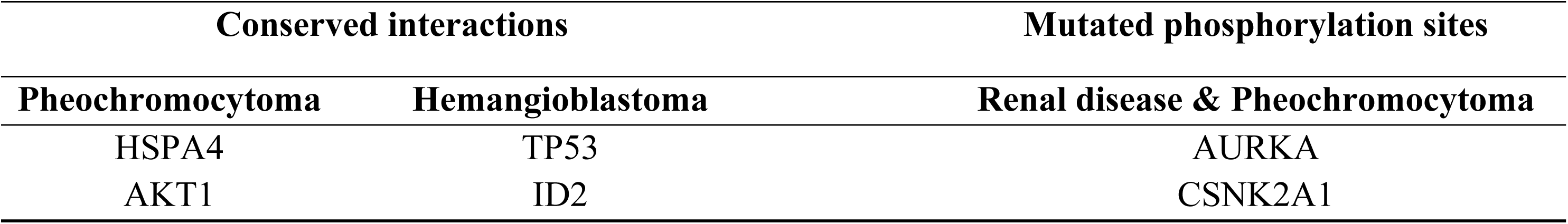
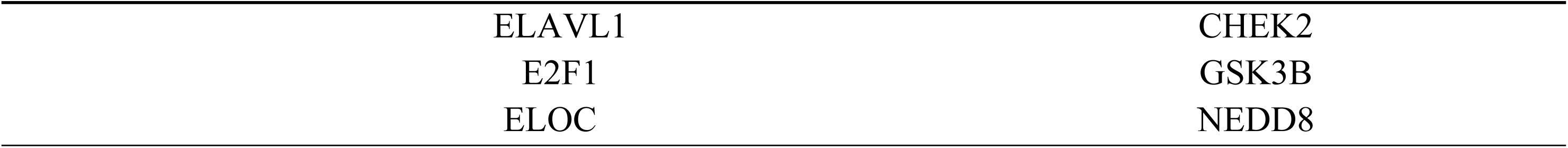
List of interactors putatively not affected by pVHL mutations found in patients presenting a specific a VHL phenotype (*left*). Phosphorylation sites found to be frequently mutated in both renal manifestations of VHL (collectively including RCC, ccRCC, renal cysts) and pheochromocytoma (*right*).

### Mutations promoting polycythemia and other interesting cases

Although a single mutation is never associated to either paraganglioma and cystadenoma as the sole phenotype observed in patients, we found specific mutations which can improve our knowledge on their onset. The five mutations p.Arg161Gln, p.Gln164His, p.Val166Phe, Arg167Trp and Arg167Gln localize in a small area forming the ELAVL1 binding interface (Fig. 5). Increased expression of this protein was recently proposed (52) to be linked with the metastatic potential of both paraganglioma and pheochromocytoma. Similarly, the three mutations p.His115Arg, p.Trp117Gly and p.Ile151Phe localize on the androgen receptor (AR) binding interface, suggesting that these specific interactors may play a crucial role in cystadenoma insurgence (Fig. 5). We also found p.Tyr175Cys linked to the insurgence of polycythemia associated with ataxia telangiectasia (AT) (53). This is a rare recessive disorder characterized by progressive cerebellar ataxia, dilatation of blood vessels in conjunctiva and eyeballs, immunodeficiency, growth retardation and sexual immaturity. Interestingly, AT is caused by mutations affecting serine-protein kinase ATM (ataxia telangiectasia mutated), a pVHL interactor (13) of unknown binding interface.

**Figure 5:**
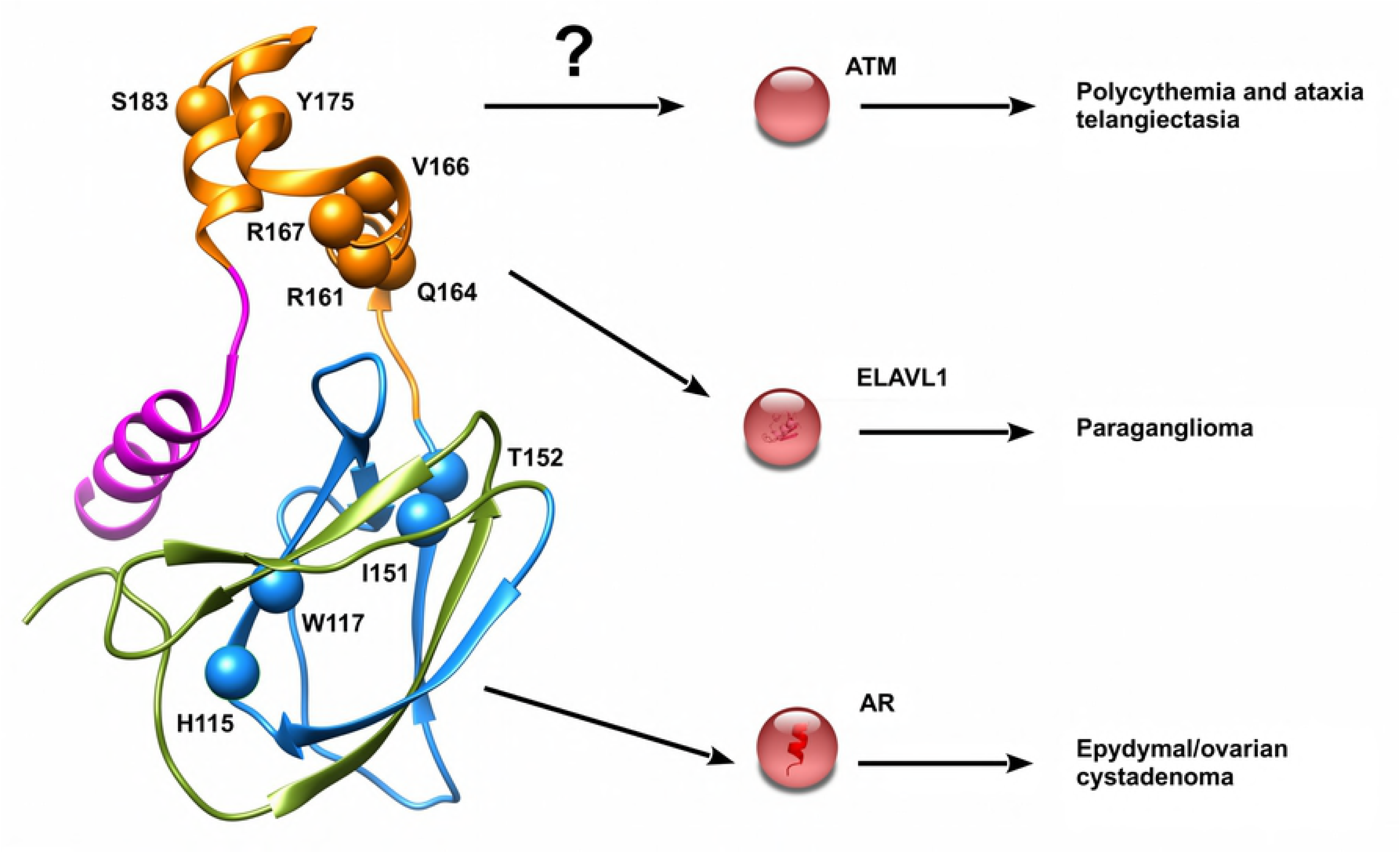
Mutations of pVHL associated to specific VHL phenotypes. Mutations promoting interesting VHL phenotypes are presented as red spheres. Interactors putatively affected and other less frequent phenotypes are reported on right.

### Predicting pVHL association with the most promising interactors

We wondered whether the binding interface data may be used to perform protein docking studies of pVHL with its multiple binding partners. We modeled the pVHL interaction with ELAVL1 and AR. Association with ELAVL1 is sustained by the pVHL α-domain and the RRM1 domain of ELAVL1 (54). As the pVHL α-domain is relevant for VCB complex formation, we first searched for possible structural analogy to perform docking by homology, but no suitable template was found. Multiple models are generated and ranked through concordance with literature data. The best docking model predicts this interaction to involve both pVHL interface A and the C-terminal tail (Figure 6A). Inspection of the interacting residues and electrostatic surface analysis shows this association to involve several electrostatic interactions. A mostly negatively charged pVHL pocket accounts for 7.2% of the pVHL accessible surface interacting with an ELAVL1 area of opposite charge (Fig. S1). Based on this model, the five mutations p.Arg161Gln, p.Gln164His, p.Val166Phe, Arg167Trp and Arg167Gln are not directly involved in ELAVL1 binding, but rather allow the right positioning of helices forming the pVHL α-domain.

**Figure 6:**
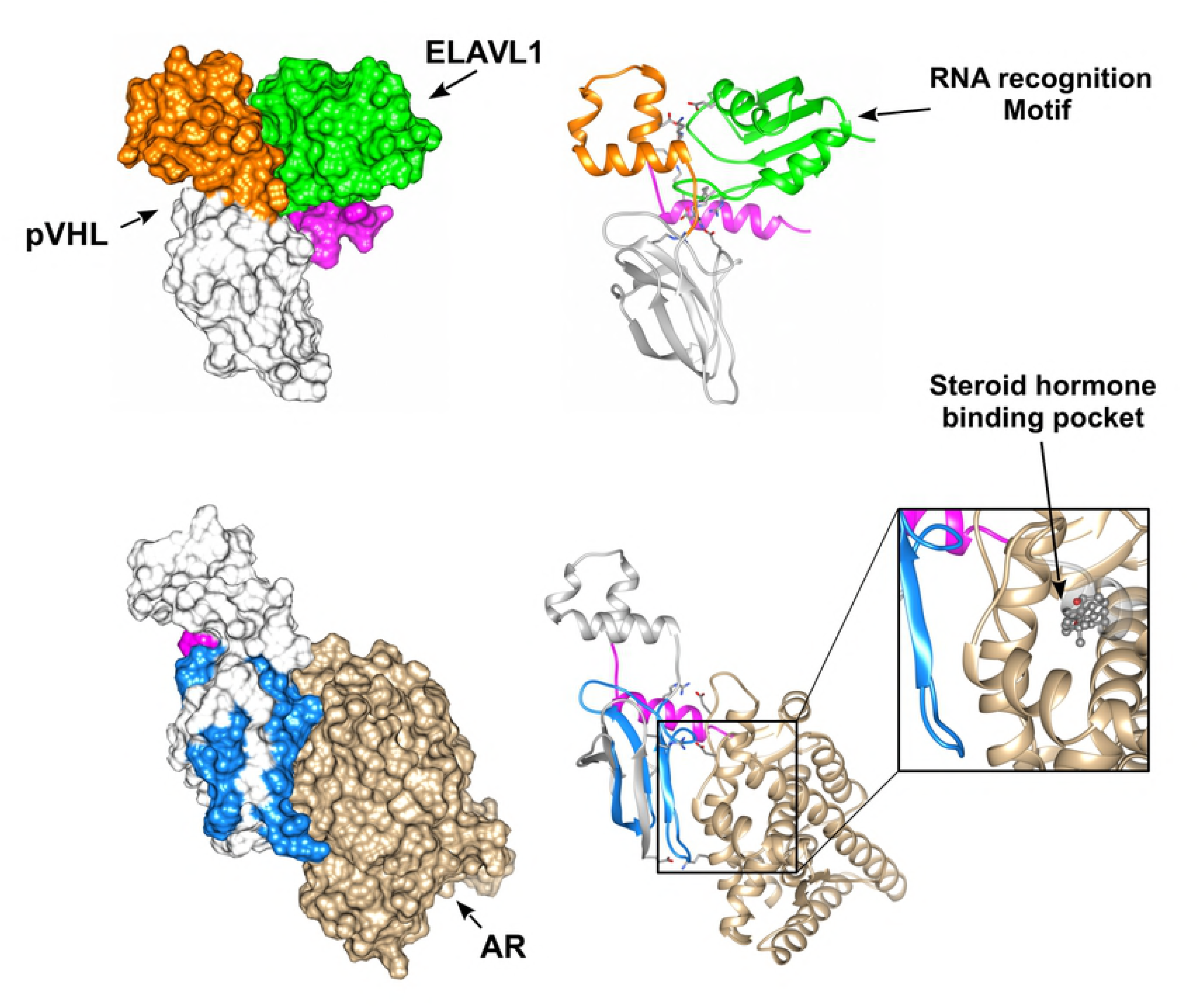
Putative pVHL molecular complexes with ELAVL1 and AR. The best scoring complexes are presented and colored according pVHL interface drawing the interaction. ELAVL1 is presented in green while brown is for AR. Zoom shows the AR ligand-binding pocket predicted to be impaired upon pVHL interaction.

The docking model of the pVHL-AR complex shows pVHL surface C interacting with α-helices H3 and H11 (Fig. 6 B) of the AR ligand-binding pocket (55). This justifies the inhibitory effect, as pVHL mutations p.His115Arg, p.Trp117Gly and p.Ile151Phe are predicted to disturb association with AR and directly interfering with correct folding of interface C. We observed that both pVHL interactions with ELAVL1 and AR are predicted as mainly electrostatic. This is coherent with the comparatively high number of exposed charged residues present in pVHL, i.e. 27.2% positive and 14.7% negative residues. On the other hand, these findings also suggest that small clusters of pVHL mutations can disturb the binding of entire pVHL interfaces by promoting local unfolding and disturbing a specific binding area. This mechanism may be easily used by cancer cells to inactivate an entire set of pVHL functions by single mutation, rapidly acquiring a fitness advantage over neighboring healthy cells.

### Simulating pVHL pathway impairments

To better clarify how cell modifications due to differentially impaired pVHL interactors prompt tumor transformation, we simulated the most promising impaired associations using a Petri net description of the pVHL interaction network (27). Considering the interesting findings from the docking analysis, we simulated the impaired associations of pVHL with ELAVL1 and AR. Interaction of pVHL with ELAVL1 is thought to play a role in p53 expression and stabilization(56). Loss of this interaction is predicted by our model to induce enhanced VEGF production and stabilization through NR4A1 (57) (Nuclear receptor subfamily 4 group A member 1) mediated inhibition of HIF-1α degradation. In parallel, we registered a glucidic metabolism imbalance due to overproduction of pro-opiomelanocortin (POMC), the precursor of the proopiomelanocortin hormone. We also observed deregulation of the Krebs cycle paired with partial PHD2 inhibition from sub-products of the carbohydrate metabolism. Alterations of proopiomelanocortin levels are a common trait of VHL-related tumors (58,59). These findings show that several mutations potentially impairing the interaction of pVHL with ELAVL1 may induce severe cell adaptations promoting tumor transformation.

AR is a steroid hormone receptor playing a pivotal role in cancer through androgen-induced gene transcription and kinase-signaling cascade activation (60,61). pVHL is thought to suppress its activity, inhibiting both genomic and nongenomic AR functions (62). Loss of pVHL-mediated AR inhibition is predicted as hyper-activating MAPK/ERK (mitogen-activated protein kinase) signaling. Under physiological conditions, MAPK/ERK activation is also stimulated by VEGF(63). We registered enhanced vascular growth and augmented vascular permeability mimicking improved VEGF production. In parallel, we observed an increased metabolic activity sustained by the large oxygen availability generated from new vessels formation. Intriguingly, proliferative pathways typical of malignant progression are predicted to be inactivated, in particular, by JADE-1 tumor-suppressive functions through induction of apoptosis (64) and Wnt signaling inhibition (65). Our model suggests a benignant phenotype characterized by high vascularization but reduced proliferation. This behavior is in agreement with the cystadenoma insurgence observed in patients harboring pVHL mutations localizing on the AR binding interface (p.His115Arg, p.Trp117Gly and p.Ile151Phe).

We also wondered whether this approach can be used to simulate loss of pVHL interaction with different kinases. We selected GSK3β and ATK-1, as representatives of phosphorylation-mediated pVHL modulation and pVHL-dependent kinase inhibition respectively. GSK3β is known to phosphorylate pVHL at Ser72 (66). This modification is linked with micro-tubule instability and external matrix remodeling. Our model predicts that upon impairment of GSK3β/pVHL association, glycogen metabolism is also impaired as enzymes deputed to glycogen synthesis are hyper-activated. Enhanced GSK3β-mediated phosphorylation of HIF-1α and its concurrent pVHL-independent degradation (67) are also predicted. In a real cell, these findings can be seen as activation of a cellular reserve mechanism for adapting its function in response to tumor suppressor mutations. GSK3β is inhibited upon insulin stimulation (68), suggesting that cells harboring mutations affecting GSK3β/pVHL association may undergo relevant HIF-1α functional deregulation when exposed to concurrent hypoxia and insulin stimuli. Finally, as pVHL is thought to inhibit AKT1, loss of pVHL association with AKT1 is predicted to promote AKT1 hyper-activation (46), which in turn promotes activation of cell survival pathways, microtubules glycolysis as well as external matrix destabilization. In a cellular environment, these modifications can collectively be interpreted as enhanced anchorage-independent growth and tumor transformation.

## Discussion

We investigated the effect of mutations affecting the pVHL tumor suppressor and their correlation with different phenotypes described for VHL patients. Our efforts in deciphering pVHL functions arise from the consideration that clinical manifestations of this familiar predisposition to develop cancers may vary among patients harboring the same mutation. The best known pVHL function is its role in degrading the HIF-1α transcription factor (8). Its role as hypoxia sensing component was conveniently used to explain some of the main VHL manifestations. However, it fails almost entirely in predicting the pathogenic risk of several mutations not directly connected with HIF-1α degradation. A robust body of literature is prompting that pVHL possesses other HIF-independent functions which co-participate in explaining the difference in disease progression clinically observed in patients (12,69,70). We previously presented VHLdb, a database collecting interactors and mutations of the human pVHL (23) aimed at rationalizing the existing knowledge around pVHL. This data is used here as a starting point to shed light on VHL disease. As a first result, we found that the most frequent mutations impair either the pVHL role in VCB complex assembly or promote β-domain destabilization. Unsurprisingly, these findings confirm that the most relevant pathogenic effect related to VHL insurgence is an inactivation of the E3-ligase function. However, an analysis of pVHL interactors putatively affected by these same mutations tells a more complex story. Our investigations hint at the possibility of associating a specific function to each pVHL surface. Based on our findings, we propose interfaces A and C of pVHL to be mainly associated with proteins involved in gene transcription and regulation and interface B to regulate protein homeostasis of several pVHL interactors. A novel C-terminal interface addresses ECM and ciliogenesis associated pVHL functions. The difference in binding property may also reflect different contributions to disease manifestations. Not all VHL patients develop the same phenotypes, in particular hemangioblastoma and renal disease and are found to be the predominant manifestations. Renal disease appears to be equally represented by mutations affecting the three main pVHL interfaces, while it is the predominant phenotype described for mutations localizing on surface D. This interface is formed by a long intrinsically disordered tail present only in the pVHL30 isoform, suggesting this specific isoform to play a precise role in renal cancer insurgence. Both paraganglioma and cystadenoma were never found as single VHL phenotypes, suggesting that these two tumors arise as secondary manifestations of VHL, possibly pairing pVHL impairment with the functional inactivation of other relevant players. We also found several mutations affecting pVHL PTM sites, indicating that malignant phenotypes can arise from pVHL functional deregulation rather than structural disruption. In particular, our simulation of phosphorylation events impaired by cancer-related mutations shows that correct interpretation of VHL fate benefits from the integration of different information sources. Manual curation and interpretation of literature data can represent a powerful tool to decipher the molecular role of this tumor suppressor protein. Collectively taken, our findings provide direct biological insights into VHL-associated tumors and may help designing novel experimental investigations to elucidate novel treatment paradigms for VHL syndrome.

## Methods

Germline and somatic pVHL mutations were manually curated from the literature from VHLdb (23), covering the most relevant literature published from 1994 to 2017 (S1 Table). Papers were included in this analysis if they contained information about the nucleotide and/or protein change (e.g. c.499C>T and/or p.Arg167Trp) as well as the pathogenic phenotype associated to mutations (e.g. RCC, CNS hemangioblastoma). A total of 1,597 different mutations from 2,123 patients were collected. pVHL surfaces were defined as previously described (22,23). Briefly, interface A spans pVHL residues 154-189, B residues 60-103 and C residues 104-153. Interface D, formed by residues 1-54 is present only in the pVHL30 isoform. Missense mutations were mapped using Chimera (71) on three different pVHL 3D-structures representing the VCB complex (PDB codes: 1LM8, 1LQB, 4WQO). As of December 2017, pVHL 3D structures cover the structured domains corresponding to both the α- and β-domains and the C-terminal region but lack the first 53 protein residues forming an intrinsically disordered N-terminal tail in pVHL30. Mutations localizing in the N-terminus region were therefore excluded from the structural analysis. Networks of interacting residues affected by mutations were calculated with RING 2.0 (28). Secondary structure and intrinsically disorder content were calculated with FELLS (72). The pVHL interactors were retrieved from VHLdb (23) (high confidence dataset). A total of 59 different interactors with known pVHL binding surface from the literature (S2 Table), were associated to pVHL mutations to highlight interactions or post-translational modification (PTM) sites are negatively affected by one or more pVHL mutations. The protein interaction network around each pVHL interface was analyzed with Cytoscape (73). VHLdb interactors were manually imported and used to investigate interactions putatively affected by mutations. Interactor clusters were investigated with MCODE (31), using VHLdb data as first shell and a second shell of STRING (74) interactors (<= 10 interactors, confidence >= 0.700, no text mining). Highly interconnected interactor regions were obtained for each pVHL interface, suggesting that these interfaces have different roles and are involved in different pathways (e.g. protein complex formation, gene transcription, signaling). These 59 interactors were in turn associated with a specific VHL disease phenotype to identify pathways impaired by pVHL missense mutations. Renal VHL manifestations (i.e. RCC, ccRCC, cysts) were collectively presented as renal disease due to many case report papers describing novel VHL mutations lacking a precise typization of cancer sub-type, e.g. generically referring to either benign renal cancer or renal carcinoma. Intrinsic disorder content was evaluated using MobiDB 3.0 (75). Molecular docking was performed using Hex(76), selecting “Shape+correlation+DARS” as correlation type and OPLS minimizations post processing. A total of 50.000 solutions were generated for each run. The PDB structures of pVHL (7) (PDB code 1LM8, chain V), human androgen receptor (55) (AR, PDB code 2AM9, chain A) and ELAVL (77) (PDB code 3H19, chain A) were retrieved from the RCSB database (78). Docking models were ranked using CONSRANK (79) and the ten best scoring models visually inspected to select models fitting literature data best, i.e. concordance between interacting interfaces and described function.

The effect of the most promising pVHL interaction impaired by the most recurrent pVHL mutations were inspected using the pVHL pathway Petri net model (27). Loss of association with AR (Human androgen receptor, UniProt ID P10275) was modeled increasing the number of tokens describing MAPK activation, i.e. p_54. Similarly, the token number of places p_38, p39 and p_60 was increased to simulate loss of pVHL-dependent AKT-1 regulation. Mutations affecting pVHL interaction with ELAVL1, GSK3β were modeled through knock-outs of the corresponding transition, i.e. t_181, t_109. For each experiment, accumulation of tokens in specific places was inspected after 2,000 simulation steps and compared with the wild-type pVHL network. A schematic representation of the analysis workflow is presented in S8 Figure.

## Acknowledgements

This work was supported by Associazione Italiana per la Ricerca sul Cancro (AIRC) grants MFAG12740 and IG17753 to ST. The funders had no role in study design, data collection and analysis, decision to publish, or preparation of the manuscript.

## Author contributions

GM, ST conceived the experiments. FQ, GM and FT performed the experiments. GM, FQ, analyzed the data. GM and ST wrote the manuscript.

## Supporting information files

**S1 Table.** Manually curated pVHL mutations. Mutations were ranked by number of affected patients.

**S2 Table.** Short description of VHL-associated cancers.

**S3 Table.** Association between mutation position and VHL manifestations. Mutations amino acid positions were associated with each corresponding pVHL surface.

**S4 Table**. Dataset of VHL mutations divided for each associated cancer.

**S5 Table.** Supplementary literature. List of scientific works used to retrieve detailed information about VHL phenotypes.

**S6 Table.** High quality pVHL interactors. List of manually curated pVHL interactors extracted from VHLdb.

**S7 File.** List of mutations affecting pVHL interactors. Interactors binding interfaces are represented as yellow blocks according with bibliographic data.

**S8 File**. **Result from MCODE analysis**. Clusters of interactors of interactors associated to each pVHL surface.

**S9 Figure.** Electrostatic surface of predicted pVHL molecular complexes. (A) Electrostatic surfaces of pVHL in complex with ELAVL1 (up) and AR (down), with red representing negatively charged areas, while blue is for positive region. Proteins are alternatively presented as transparent views and rotated around vertical axis to better highlight complementary surfaces.

**S10 Figure.** Schematic representation of the analysis workflow.

## References

1. Hemminki K, Vaittinen P. Familial cancers in a nationwide family cancer database: age distribution and prevalence. Eur J Cancer. 1999 Jul;35(7):1109–17.

2. Stiller CA. Epidemiology and genetics of childhood cancer. Oncogene. 2004;23(38):6429–44.

3. Scheuner MT, McNeel TS, Freedman AN. Population prevalence of familial cancer and common hereditary cancer syndromes. The 2005 California Health Interview Survey. Genet Med. 2010 Nov;12(11):726–35.

4. Iliopoulos O, Kibel A, Gray S, Kaelin WG. Tumour suppression by the human von Hippel-Lindau gene product. Nat Med. 1995 Agosto;1(8):822–6.

5. Barry RE, Krek W. The von Hippel-Lindau tumour suppressor: a multi-faceted inhibitor of tumourigenesis. Trends in molecular medicine. 2004 Sep;10(9):466–72.

6. Keutgen XM, Hammel P, Choyke PL, Libutti SK, Jonasch E, Kebebew E. Evaluation and management of pancreatic lesions in patients with von Hippel–Lindau disease. Nature Reviews Clinical Oncology. 2016 Sep;13(9):537–49.

7. Min J-H, Yang H, Ivan M, Gertler F, Kaelin WG, Pavletich NP. Structure of an HIF-1alpha-pVHL complex: hydroxyproline recognition in signaling. Science. 2002 Jun 7;296(5574):1886–9.

8. Kim WY, Kaelin WG. Role of VHL gene mutation in human cancer. J Clin Oncol. 2004 Dec 15;22(24):4991–5004.

9. Berra E, Benizri E, Ginouvès A, Volmat V, Roux D, Pouysségur J. HIF prolyl-hydroxylase 2 is the key oxygen sensor setting low steady-state levels of HIF-1α in normoxia. The EMBO Journal. 2003 Aug 15;22(16):4082–90.

10. Minervini G, Quaglia F, Tosatto SCE. Insights into the proline hydroxylase (PHD) family, molecular evolution and its impact on human health. Biochimie. 2015 Settembre;116:114–24.

11. Nordstrom-O’Brien M, van der Luijt RB, van Rooijen E, van den Ouweland AM, Majoor-Krakauer DF, Lolkema MP, et al. Genetic analysis of von Hippel-Lindau disease. Hum Mutat. 2010 May 1;31(5):521–37.

12. Crespigio J, Berbel LCL, Dias MA, Berbel RF, Pereira SS, Pignatelli D, et al. Von Hippel–Lindau disease: a single gene, several hereditary tumors. J Endocrinol Invest. 2017 Jun 6;1–11.

13. Roe J-S, Kim H, Lee S-M, Kim S-T, Cho E-J, Youn H-D. p53 Stabilization and Transactivation by a von Hippel-Lindau Protein. Molecular Cell. 2006 Maggio;22(3):395–405.

14. Clifford SC, Cockman ME, Smallwood AC, Mole DR, Woodward ER, Maxwell PH, et al. Contrasting effects on HIF-1alpha regulation by disease-causing pVHL mutations correlate with patterns of tumourigenesis in von Hippel-Lindau disease. Hum Mol Genet. 2001 May 1;10(10):1029–38.

15. Grosfeld A, Stolze IP, Cockman ME, Pugh CW, Edelmann M, Kessler B, et al. Interaction of hydroxylated collagen IV with the von hippel-lindau tumor suppressor. J Biol Chem. 2007 May 4;282(18):13264–9.

16. Kurban G, Duplan E, Ramlal N, Hudon V, Sado Y, Ninomiya Y, et al. Collagen matrix assembly is driven by the interaction of von Hippel-Lindau tumor suppressor protein with hydroxylated collagen IV alpha 2. Oncogene. 2008 Feb 7;27(7):1004–12.

17. Ji Q, Burk RD. Downregulation of integrins by von Hippel-Lindau (VHL) tumor suppressor protein is independent of VHL-directed hypoxia-inducible factor alpha degradation. Biochem Cell Biol. 2008 Jun;86(3):227–34.

18. Young AP, Schlisio S, Minamishima YA, Zhang Q, Li L, Grisanzio C, et al. VHL loss actuates a HIF-independent senescence programme mediated by Rb and p400. Nat Cell Biol. 2008 Mar;10(3):361–9.

19. Lee S, Nakamura E, Yang H, Wei W, Linggi MS, Sajan MP, et al. Neuronal apoptosis linked to EglN3 prolyl hydroxylase and familial pheochromocytoma genes: developmental culling and cancer. Cancer Cell. 2005 Aug;8(2):155–67.

20. Yang H, Minamishima YA, Yan Q, Schlisio S, Ebert BL, Zhang X, et al. pVHL acts as an adaptor to promote the inhibitory phosphorylation of the NF-kappaB agonist Card9 by CK2. Mol Cell. 2007 Oct 12;28(1):15–27.

21. Kuznetsova AV, Meller J, Schnell PO, Nash JA, Ignacak ML, Sanchez Y, et al. von Hippel-Lindau protein binds hyperphosphorylated large subunit of RNA polymerase II through a proline hydroxylation motif and targets it for ubiquitination. Proc Natl Acad Sci USA. 2003 Mar 4;100(5):2706–11.

22. Leonardi E, Murgia a, Tosatto SCE. Adding structural information to the von Hippel-Lindau (VHL) tumor suppressor interaction network. FEBS letters. 2009 Nov;583(22):3704–10.

23. Tabaro F, Minervini G, Sundus F, Quaglia F, Leonardi E, Piovesan D, et al. VHLdb: A database of von Hippel-Lindau protein interactors and mutations. Scientific Reports. 2016 Aug 11;6:31128.

24. Esteban-Barragán MA, Avila P, Alvarez-Tejado M, Gutiérrez MD, García-Pardo A, Sánchez-Madrid F, et al. Role of the von Hippel-Lindau tumor suppressor gene in the formation of beta1-integrin fibrillar adhesions. Cancer Res. 2002 May 15;62(10):2929–36.

25. Li M, Kim WY. Two sides to every story: the HIF-dependent and HIF-independent functions of pVHL. J Cell Mol Med. 2011 Feb;15(2):187–95.

26. Nguyen HC, Yang H, Fribourgh JL, Wolfe LS, Xiong Y. Insights into Cullin-RING E3 Ubiquitin Ligase Recruitment: Structure of the VHL-EloBC-Cul2 Complex. Structure [Internet]. [cited 2015 Feb 13]; Available from: http://www.sciencedirect.com/science/article/pii/S0969212614004237

27. Minervini G, Panizzoni E, Giollo M, Masiero A, Ferrari C, Tosatto SCE. Design and analysis of a Petri net model of the Von Hippel-Lindau (VHL) tumor suppressor interaction network. PLoS ONE. 2014;9(6):e96986.

28. Piovesan D, Minervini G, Tosatto SCE. The RING 2.0 web server for high quality residue interaction networks. Nucleic Acids Res. 2016 May 19;

29. Robinson CM, Ohh M. The multifaceted von Hippel-Lindau tumour suppressor protein. FEBS Lett. 2014 Aug 19;588(16):2704–11.

30. Roe J-S, Youn H-D. The Positive Regulation of p53 by the Tumor Suppressor VHL. Cell Cycle. 2006 Sep 15;5(18):2054–6.

31. Bader GD, Hogue CW. An automated method for finding molecular complexes in large protein interaction networks. BMC Bioinformatics. 2003 Jan 13;4:2.

32. Vainberg IE, Lewis SA, Rommelaere H, Ampe C, Vandekerckhove J, Klein HL, et al. Prefoldin, a chaperone that delivers unfolded proteins to cytosolic chaperonin. Cell. 1998 May 29;93(5):863–73.

33. Russell RC, Sufan RI, Zhou B, Heir P, Bunda S, Sybingco SS, et al. LOSS OF JAK2 REGULATION VIA VHL-SOCS1 E3 UBIQUITIN HETEROCOMPLEX UNDERLIES CHUVASH POLYCYTHEMIA. Nat Med. 2011 Jun 19;17(7):845–53.

34. Berthouze M, Venkataramanan V, Li Y, Shenoy SK. The deubiquitinases USP33 and USP20 coordinate beta2 adrenergic receptor recycling and resensitization. EMBO J. 2009 Jun 17;28(12):1684–96.

35. Mahon PC, Hirota K, Semenza GL. FIH-1: a novel protein that interacts with HIF-1α and VHL to mediate repression of HIF-1 transcriptional activity. Genes Dev. 2001 Oct 15;15(20):2675–86.

36. Zhao W-T, Zhou C-F, Li X-B, Zhang Y-F, Fan L, Pelletier J, et al. The von Hippel-Lindau protein pVHL inhibits ribosome biogenesis and protein synthesis. J Biol Chem. 2013 Jun 7;288(23):16588–97.

37. Anderson K, Nordquist KA, Gao X, Hicks KC, Zhai B, Gygi SP, et al. Regulation of cellular levels of Sprouty2 protein by prolyl hydroxylase domain and von Hippel-Lindau proteins. J Biol Chem. 2011 Dec 9;286(49):42027–36.

38. Xie L, Xiao K, Whalen EJ, Forrester MT, Freeman RS, Fong G, et al. Oxygen-regulated beta(2)-adrenergic receptor hydroxylation by EGLN3 and ubiquitylation by pVHL. Sci Signal. 2009 Jul 7;2(78):ra33.

39. Heir P, Srikumar T, Bikopoulos G, Bunda S, Poon BP, Lee JE, et al. Oxygen-dependent Regulation of Erythropoietin Receptor Turnover and Signaling. J Biol Chem. 2016 Apr 1;291(14):7357–72.

40. Jin H, White SR, Shida T, Schulz S, Aguiar M, Gygi SP, et al. The conserved Bardet-Biedl syndrome proteins assemble a coat that traffics membrane proteins to cilia. Cell. 2010 Jun 25;141(7):1208–19.

41. Minervini G, Mazzotta GM, Masiero A, Sartori E, Corrà S, Potenza E, et al. Isoform-specific interactions of the von Hippel-Lindau tumor suppressor protein. Sci Rep. 2015;5:12605.

42. Lai Y, Song M, Hakala K, Weintraub ST, Shiio Y. Proteomic dissection of the von Hippel-Lindau (VHL) interactome. J Proteome Res. 2011 Nov 4;10(11):5175–82.

43. Frew IJ, Krek W. pVHL: a multipurpose adaptor protein. Sci Signal. 2008;1(24):pe30.

44. Russell RC, Ohh M. NEDD8 acts as a ‘molecular switch’’ defining the functional selectivity of VHL.’ EMBO Rep. 2008 May;9(5):486–91.

45. Carpten JD, Faber AL, Horn C, Donoho GP, Briggs SL, Robbins CM, et al. A transforming mutation in the pleckstrin homology domain of AKT1 in cancer. Nature. 2007 Jul 26;448(7152):439–44.

46. Guo J, Chakraborty AA, Liu P, Gan W, Zheng X, Inuzuka H, et al. pVHL suppresses kinase activity of Akt in a proline-hydroxylation–dependent manner. Science. 2016 Aug 26;353(6302):929–32.

47. Dyer KD, Lavigne MC, Rosenberg HF. Hsp70RY: further characterization of a novel member of the hsp70 protein family. Biochem Biophys Res Commun. 1994 Aug 30;203(1):577–81.

48. Yang C, Huntoon K, Ksendzovsky A, Zhuang Z, Lonser RR. Proteostasis modulators prolong missense VHL protein activity and halt tumor progression. Cell Rep. 2013 Jan 31;3(1):52–9.

49. Hescot S, Leboulleux S, Amar L, Vezzosi D, Borget I, Bournaud-Salinas C, et al. One-year progression-free survival of therapy-naive patients with malignant pheochromocytoma and paraganglioma. J Clin Endocrinol Metab. 2013 Oct;98(10):4006–12.

50. Na X, Duan HO, Messing EM, Schoen SR, Ryan CK, di Sant‘Agnese PA, et al. Identification of the RNA polymerase II subunit hsRPB7 as a novel target of the von Hippel-Lindau protein. EMBO J. 2003 Aug 15;22(16):4249–59.

51. Mikhaylova O, Ignacak ML, Barankiewicz TJ, Harbaugh SV, Yi Y, Maxwell PH, et al. The von Hippel-Lindau tumor suppressor protein and Egl-9-Type proline hydroxylases regulate the large subunit of RNA polymerase II in response to oxidative stress. Mol Cell Biol. 2008 Apr;28(8):2701–17.

52. Leijon H, Salmenkivi K, Heiskanen I, Hagström J, Louhimo J, Heikkilä P, et al. HuR in pheochromocytomas and paragangliomas - overexpression in verified malignant tumors. APMIS. 2016 Sep;124(9):757–63.

53. Bento MC, Chang KT, Guan Y, Liu E, Caldas G, Gatti RA, et al. Congenital polycythemia with homozygous and heterozygous mutations of von Hippel-Lindau gene: five new Caucasian patients. Haematologica. 2005 Jan;90(1):128–9.

54. Datta K, Mondal S, Sinha S, Li J, Wang E, Knebelmann B, et al. Role of elongin-binding domain of von hippel lindau gene product on HuR-mediated VPF/VEGF mRNA stability in renal cell carcinoma. Oncogene. 2005 Nov;24(53):7850.

55. Pereira de Jésus-Tran K, Côté P-L, Cantin L, Blanchet J, Labrie F, Breton R. Comparison of crystal structures of human androgen receptor ligand-binding domain complexed with various agonists reveals molecular determinants responsible for binding affinity. Protein Science. 2006 May 1;15(5):987–99.

56. Galba S, Martindale JL, Mazan-mamczarz K, Lo I, Fan J, Wang W, et al. Influence of the RNA-Binding Protein HuR in pVHL-Regulated p53 Expression in Renal Carcinoma Cells. 2003;23(20):7083–7095.

57. Kim BY, Kim H, Cho EJ, Youn HD. Nur77 upregulates HIF-alpha by inhibiting pVHL-mediated degradation. Exp Mol Med. 2008 Feb 29;40(1):71–83.

58. DeBold CR, Mufson EE, Menefee JK, Orth DN. Proopiomelanocortin gene expression in a pheochromocytoma using upstream transcription initiation sites. Biochem Biophys Res Commun. 1988 Sep 15;155(2):895–900.

59. Choi J-W, Park SC, Kang GH, Liu JO, Youn H-D. Nur77 activated by hypoxia-inducible factor-1alpha overproduces proopiomelanocortin in von Hippel-Lindau-mutated renal cell carcinoma. Cancer Res. 2004 Jan 1;64(1):35–9.

60. Heinlein CA, Chang C. The Roles of Androgen Receptors and Androgen-Binding Proteins in Nongenomic Androgen Actions. Mol Endocrinol. 2002 Oct 1;16(10):2181–7.

61. Chen CD, Welsbie DS, Tran C, Baek SH, Chen R, Vessella R, et al. Molecular determinants of resistance to antiandrogen therapy. Nat Med. 2004 Jan;10(1):33–9.

62. Wang J, Zhang W, Ji W, Liu X, Ouyang G, Xiao W. The Von Hippel-Lindau Protein Suppresses Androgen Receptor Activity. Mol Endocrinol. 2014 Feb 1;28(2):239–48.

63. Doanes AM, Hegland DD, Sethi R, Kovesdi I, Bruder JT, Finkel T. VEGF stimulates MAPK through a pathway that is unique for receptor tyrosine kinases. Biochem Biophys Res Commun. 1999 Feb 16;255(2):545–8.

64. Zhou MI, Foy RL, Chitalia VC, Zhao J, Panchenko MV, Wang H, et al. Jade-1, a candidate renal tumor suppressor that promotes apoptosis [Internet]. [cited 2016 Jan 12]. Available from: http://www.pnas.org

65. Chitalia VC, Foy RL, Bachschmid MM, Zeng L, Panchenko MV, Zhou MI, et al. Jade-1 inhibits Wnt signalling by ubiquitylating beta-catenin and mediates Wnt pathway inhibition by pVHL. Nat Cell Biol. 2008 Oct;10(10):1208–16.

66. Hergovich A, Lisztwan J, Thoma CR, Wirbelauer C, Barry RE, Krek W. Priming-Dependent Phosphorylation and Regulation of the Tumor Suppressor pVHL by Glycogen Synthase Kinase 3. Mol Cell Biol. 2006 Aug;26(15):5784–96.

67. Flügel D, Görlach A, Michiels C, Kietzmann T. Glycogen synthase kinase 3 phosphorylates hypoxia-inducible factor 1alpha and mediates its destabilization in a VHL-independent manner. Mol Cell Biol. 2007 May;27(9):3253–65.

68. Cross DA, Alessi DR, Cohen P, Andjelkovich M, Hemmings BA. Inhibition of glycogen synthase kinase-3 by insulin mediated by protein kinase B. Nature. 1995 Dec 21;378(6559):785–9.

69. Gossage L, Eisen T, Maher ER. VHL, the story of a tumour suppressor gene. Nat Rev Cancer. 2015 Gennaio;15(1):55–64.

70. Tarade D, Ohh M. The HIF and other quandaries in VHL disease. Oncogene [Internet]. 2017 Sep 18 [cited 2017 Sep 21]; Available from: http://www.nature.com/onc/journal/vaop/ncurrent/abs/onc2017338a.html?foxtrotcallback=true

71. Pettersen EF, Goddard TD, Huang CC, Couch GS, Greenblatt DM, Meng EC, et al. UCSF Chimera a visualization system for exploratory research and analysis. J Comput Chem. 2004 Oct;25(13):1605–12.

72. Piovesan D, Walsh I, Minervini G, Tosatto SCE. FELLS: fast estimator of latent local structure. Bioinformatics. 2017 Feb 10;

73. Shannon P, Markiel A, Ozier O, Baliga NS, Wang JT, Ramage D, et al. Cytoscape: A Software Environment for Integrated Models of Biomolecular Interaction Networks. Genome Res. 2003 Nov 1;13(11):2498–504.

74. Szklarczyk D, Morris JH, Cook H, Kuhn M, Wyder S, Simonovic M, et al. The STRING database in 2017: quality-controlled protein–protein association networks, made broadly accessible. Nucleic Acids Res. 2017 Jan 4;45(Database issue):D362–8.

75. Piovesan D, Tabaro F, Paladin L, Necci M, Micetic I, Camilloni C, et al. MobiDB 3.0: more annotations for intrinsic disorder, conformational diversity and interactions in proteins. Nucleic Acids Res [Internet]. [cited 2017 Dec 4]; Available from: https://academic.oup.com/nar/advance-article/doi/10.1093/nar/gkx1071/4612964

76. Ritchie DW, Venkatraman V. Ultra-fast FFT protein docking on graphics processors. Bioinformatics. 2010 Oct 1;26(19):2398–405.

77. Benoit RM, Meisner N-C, Kallen J, Graff P, Hemmig R, Cèbe R, et al. The x-ray crystal structure of the first RNA recognition motif and site-directed mutagenesis suggest a possible HuR redox sensing mechanism. J Mol Biol. 2010 Apr 16;397(5):1231–44.

78. Rose PW, Prlić A, Altunkaya A, Bi C, Bradley AR, Christie CH, et al. The RCSB protein data bank: integrative view of protein, gene and 3D structural information. Nucleic Acids Res. 2017 Jan 4;45(D1):D271–81.

79. Chermak E, Petta A, Serra L, Vangone A, Scarano V, Cavallo L, et al. CONSRANK: a server for the analysis, comparison and ranking of docking models based on inter-residue contacts. Bioinformatics. 2015 May 1;31(9):1481–3.

